# Accurate estimation of neural population dynamics without spike sorting

**DOI:** 10.1101/229252

**Authors:** Eric M. Trautmann, Sergey D. Stavisky, Subhaneil Lahiri, Katherine C. Ames, Matthew T. Kaufman, Stephen I. Ryu, Surya Ganguli, Krishna V. Shenoy

## Abstract

A central goal of systems neuroscience is to relate an organism’s neural activity to behavior. Neural population analysis often begins by reducing the dimensionality of the data to focus on the patterns most relevant to a given task. A major practical hurdle to data analysis is spike sorting, and this problem is growing rapidly as the number of neurons measured increases. Here, we investigate whether spike sorting is necessary to estimate neural dynamics. The theory of random projections suggests that we can accurately estimate the geometry of low-dimensional manifolds from a small number of linear projections of the data. We re-analyzed data from three previous studies and found that neural dynamics and scientific conclusions are quite similar using multi-unit threshold crossings in place of sorted neurons. This finding unlocks existing data for new analyses and informs the design and use of new electrode arrays for laboratory and clinical use.

## Introduction

A growing number of studies, spanning systems neuroscience, seek to relate the dynamical evolution of neural population states with an organism’s behavior (e.g., Briggman et al. (2005); Machens et al. (2010); Harvey et al. (2012); Churchland et al. (2012); Mante et al. (2013); Ames et al. (2014); Sadtler et al. (2014); Kaufman et al. (2014, 2015); Morcos & Harvey (2016)). In this work, we aim to address a major challenge facing neurophysiological experiments: how can we cope with the challenge of attributing action potentials to individual neurons, termed spike sorting, as the number of electrodes rapidly increases from several hundred (today’s state of the art) to thousands or even millions in the near future Stevenson & Kording (2011)? This exciting and rapid increase is necessary for advancing neuroscientific understanding and brain-machine interfaces, and progress is fueled in part by the U.S. BRAIN initiative and similar efforts around the world Bargmann et al. (2014). For a typical experiment, composed of several hours of neural and behavioral recordings, manually spike sorting even 100 channels can take a skilled researcher several hours, and different human experts often arrive at different results Wood et al. (2004). Automated spike sorting algorithms show promise, e.g.: Santhanam et al. (2004); Vargas-Irwin & Donoghue (2007); Wood & Black (2008); Ventura (2009); Chah et al. (2011); Bestel et al. (2012); Barnett et al. (2016)), but are computationally intensive, sensitive to changes in waveform due to electrode drift, and no ground truth is available. Exciting recent methods leverage high-density neural recordings to yield reliable single neuron isolation, but such sensors represent a minority of emerging technologies as they are optimized specifically for high-density recording within a small volume around a linear probe Harris et al. (2016); Rossant et al. (2016); Pachitariu et al. (2016); Leibig et al. (2016);Jun et al. (2017a); Chung et al. (2017). This requirement creates a trade off between spike sorting quality and measuring from a larger volume of tissue. This is in contrast to the need to also measure from many regions of the brain simultaneously and to do so for long durations (i.e., chronic implants Chestek et al. (2011); Barrese et al. (2013)).

Here we investigate whether spike sorting is a necessary data pre-processing step for data analyses that focus on neural population activity. For investigations involving the response of a single neuron, or when the nature of the question requires certainty about neuron identity, spike sorting is required. However, for investigations involving the coordinated response and evolution of large populations of neurons, we ask if spike sorting is essential. In other words, does combining neurons by not spike sorting result in distorted estimates of neural population states and neural dynamics, thereby changing the results of hypothesis tests?

Brain-machine interfaces (BMIs) that measure motor cortical activity and aim to help people with paralysis have moved away from spike sorting in recent years (e.g., Gilja et al. (2015); Pandarinath et al. (2017a)). This was motivated by the considerations described above, as well as by the repeated finding that the performance difference between spike sorting and not spike sorting is quite small Chestek et al. (2011); Christie et al. (2015); Todorova et al. (2014). Most pre-clinical and clinical-trial BMIs do not currently employ spike sorting, and instead use a simple voltage threshold crossing rate, which combines the responses from all action potentials on an electrode regardless of their source neuron Fraser et al. (2009); Gilja et al. (2012); Hochberg et al. (2012); Collinger et al. (2013); Jarosiewicz et al. (2015); Gilja et al. (2015); Perel et al. (2015); Christie et al. (2015); Kao et al. (2016); Pandarinath et al. (2017a); Ajiboye et al. (2017), and alleviating the burden of spike sorting, while maintaining their high level of single-trial neural population decoding. Here we ask whether this simple and efficient threshold-based approach can be applied in basic neuroscience investigations (i.e., assessing hypotheses based on identifying structure and dynamics in neural data), where the need for spike sorting could potentially be more stringent.

In a standard extra-cellular electrophysiology experiment, a multi-electrode array gives researchers access to a sparse sample of a few hundred neurons, selected randomly from the many millions of neurons in a particular brain region. For most typical experiments, spikes are sorted to associate action potentials from individual neurons, prior to subsequent analysis steps, such as performing dimensionality reduction. The process of spike sorting expands the dimensionality of the dataset from the number of recording channels to the total number of isolatable neurons observed on the array (see Figure 1A). Here, we propose to bypass this sorting step prior to population-level analyses. Dimensionality reduction methods, such as PCA, GPFA, dPCA, or LFADS, typically use linear combinations of individual sorted neurons to capture important aspects of the population response in a reduced representation of the data (e.g., Yu et al. (2009); Churchland et al. (2012); Kobak et al. (2016); Sussillo et al. (2016); Pandarinath et al. (2017b), Figure 1B). By starting with multi-unit threshold crossings, the multiple neurons present on each channel are linearly summed prior to performing a second linear operation via dimensionality reduction, the resulting population activity closely resemble those found with sorted neurons, shown schematically in Figure 1C. Applying this to neural data from rhesus monkeys, performing radial reaching tasks, yields substantial qualitative similarity between neural trajectories using sorted units and multi-unit threshold crossings (Figure 1D-E). While these qualitative findings are suggestive, a quantitative study is required before it is possible to understand its role in neuroscientific studies. That is what we investigate in this study.

**Figure 1:**
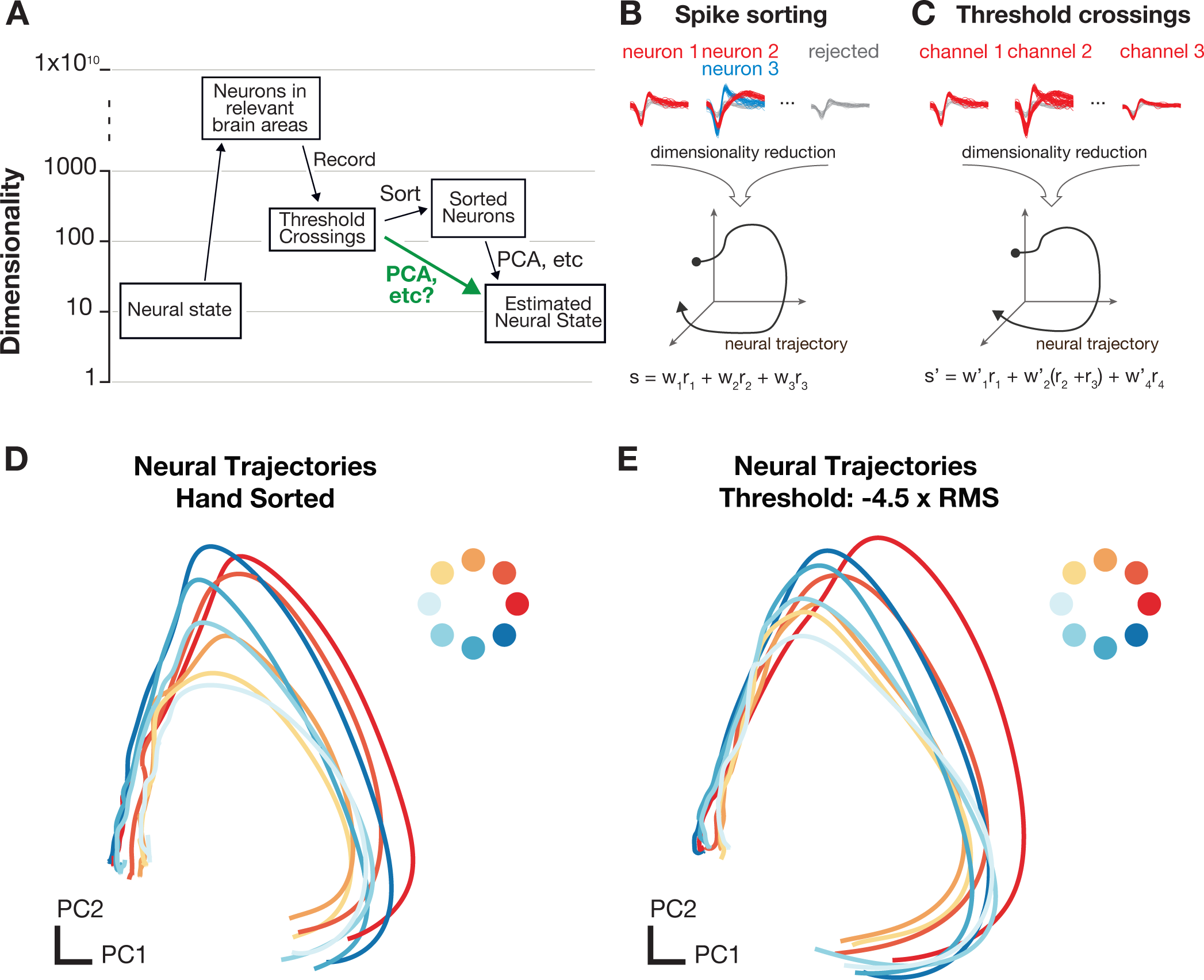
SortFree Concept. **A)** Data acquisition and preprocessing pipeline The experimentally measured dimensionality of neural activity in the motor system suggests that a small number of latent factors (often 8-12) captures the majority of task-relevant neural variability. In most experiments, neural recordings are sparsely sampled from up to a few hundred neurons in systems containing many millions of neurons. **B,C)** Typically, spikes from individual neurons are sorted using action potential waveforms and dimensionality reduction is performed on the smoothed firing rates of the isolated units. Here, we propose that for certain specific classes of analyses, it’s theoretically sensible to bypass the sorting step and perform dimensionality reduction or population-level analyses on voltage threshold crossings directly. Neural trajectories for delayed reaches to one of eight radial targets using manually sorted neurons **D)** or multi-unit threshold crossings **E)** display little distortion in the low-dimensional projections using PCA.

This approach is motivated not only by the success of BMI threshold crossing decoders, but more fundamentally by the theory of random projections from high dimensional statistics (e.g., Indyk & Motwani (1998); Dasgupta & Gupta (2003); Ganguli & Sompolinsky (2012); Advani et al. (2013); Lahiri et al. (2016)). Recent work has shown that if one wishes to recover the geometry of a low dimensional manifold, which is embedded in a high dimensional space, one can still accurately recover this geometry without measuring all of the coordinates of the high dimensional space. Instead, it is sufficient to measure a small number of noisy, random linear combinations (i.e., projections) of these coordinates Ganguli & Sompolinsky (2012). In the neuroscience context, (1) the low dimensional manifold is a surface containing the set of low dimensional neural population trajectories; (2) the coordinates in high dimensional space are the firing rates or spike counts of individual neurons; and (3) the noisy projections of these coordinates are the activities measured on each electrode, which consist of the linear combination of a small number of neurons as well as multi-unit ‘hash’ that cannot be resolved into single neuron spikes. We posit that this application of random projection theory to neural measurements suggests that spike-sorting may not be necessary to accurately recover neural population dynamics, which are inherently low dimensional.

Encouraged by recent BMI results and the theory of random projections, we replicated analyses from three previously published studies of nonhuman primate motor cortical control of arm movements, now using multi-unit threshold crossings (i.e., the linear combination of a small number of neurons as well as multi-unit ‘hash’). We compared the resulting neural population state dynamics and hypothesis outcomes with the original studies and found that all of the new analyses using multi-unit threshold crossings closely recapitulated both qualitative and quantitative features found in the original studies using spike sorted data and yielded the same scientific conclusions. We further show that the similarity of neural population dynamics extracted from sorted versus unsorted data is consistent with the theory of random projections, and we derive quantitative scaling laws for how this similarity depends on the complexity of the population dynamics themselves.

We suggest that these findings may well: (1) unlock large repositories of existing data for new analyses without time-consuming manual sorting or error-prone automatic sorting, (2) inform the design and use of new wireless, low power electrode arrays for laboratory investigations and clinical use (in particular, chronically-implanted multielectrode arrays and bandwidth and power limited wireless telemetry systems), and (3) enable scientific measurements using electrode arrays that do not afford high quality spike sorting.

## Results

To investigate the necessity of spike-sorting, we re-analyzed data collected in three recently published studies by Ames and colleages, Churchland and colleagues and Kaufman and colleagues Ames et al. (2014); Churchland et al. (2012); Kaufman et al. (2014). All three studies relate the spiking activity of populations of single neurons in macaque motor and dorsal premotor cortex to arm reaching behavior. Here we substitute multi-unit threshold crossings prior to performing the same analyses used in the original studies. As described below and in Methods, using a sufficiently large voltage threshold effectively rejects noise arising from electrical artifacts and noise. Thus, multi-unit threshold crossings predominantly correspond to action potential emission from one or multiple neurons relatively close to the electrode. See Figure S1 for a comparison of spike waveforms from sorted single units and multi-unit threshold crossings. As anticipated, the tuning for single recording channels tends to broaden (Figure S1D) and peri-movement firing rates increase (Figure S1E) as the threshold becomes more permissive.

### Study 1: Neural dynamics of reaching following incorrect or absent motor preparation

Ames and colleagues asked whether the preparatory neural population state achieved by motor cortex prior to the initiation of movement is obligatory for generating an accurate reach, either when no time is given to prepare the reach (no delay period) or when the target location switches at the time of the go cue Ames et al. (2014). The key result of the study, found using manually spike-sorted data, was that in both cases the neural population state can bypass the preparatory state when no time is provided to prepare the movement. Now, using multi-unit threshold crossing data, we observed the same results for both behavior types using the same analyses and statistical tests (*p* < 0.05; distance metric statistical test Ames et al. (2014)). In addition, visualizations of neural trajectories using the top two principal components (Figure 2A,B) reveal quite similar features (which are inherently assessed qualitatively) when using multi-unit threshold crossings instead of sorted neurons. This suggests that using multi-unit threshold crossings does not substantially distort our lower-dimensional view of neural population dynamics.

**Figure 2:**
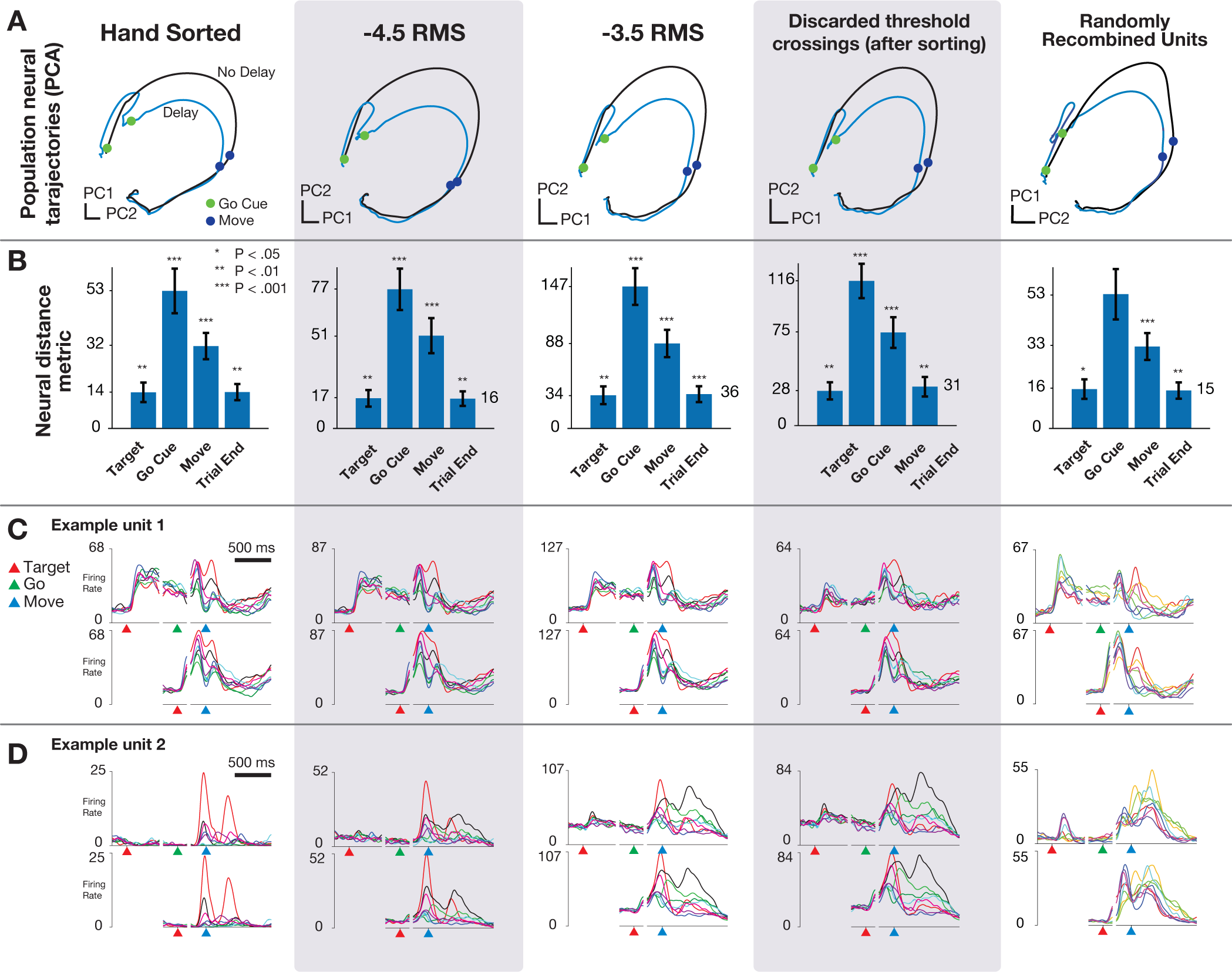
Replication of “Neural population dynamics during reaching Ames et al. (2014)”. **(A)** Neural trajectories calculated using PCA on trial-averaged neural activity for reaches with and without delay period using hand sorted units, more conservative threshold set at −4.5 × RMS, more permissive threshold set at −3.5 × RMS, using only threshold crossings that were discarded after sorting, and when sorted units were randomly combined to simulate multi-unit channels. **(B)** Key results from Ames et al. (2014), distance in full-dimensional neural space between trial-averaged neural trajectories of reaches when the monkey was or was not presented with a delay period. Note that the y-axis is scaled between columns, illustrating that although ensemble firing rates are higher with more permissive thresholds, the key qualitative and quantitative features of the population neural response are conserved. **(C)** Example unit 1: PSTHs for center-out reaches to eight radially spaced targets. In this example, the Y-axis scales upward with a more permissive threshold, but the overall shape of the PSTH is quite similar regardless of sorting or thresholding. **(D)** Example unit 2: Features of the PSTHs for this unit do change as a more permissive threshold is used. Despite this variation, the estimated neural state from the population response, as shown in **A** is largely invariant to choice of threshold.

We repeated these analyses with two simulated perturbations to the dataset to further test the sensitivity of population analyses to combining the contributions of single neurons. First, we analyzed only the multi-unit threshold crossing events that were previously discarded through spike sorting. The computed neural trajectories, distance plots, and peri-stimulus time histograms (PSTHs) for these ‘discarded spikes’ suggest that there is similar information content in these low-amplitude spikes as there are in the sorted individual neurons (Figure 2, fourth column). Second, we repeated this analysis with data created by randomly recombining individual neurons to simulate multi-unit threshold crossings. In this case, the results also closely matched those found with sorted single neurons (Figure 2, fifth column). Note that the PSTHs for this case are not anticipated to resemble those of any particular single units, but are not shown to demonstrate that recombination of units does not substantially wash out tuning.

These results afford several observations. First, for some electrode channels, the shape of the PSTH across conditions is closely recapitulated regardless of the threshold level or inclusion of hash (Figure 2C). For these channels, the spatial and temporal properties of the constituent parts of the multi-unit threshold crossings (i.e., individual neurons and hash) are similar enough that combining these components just results in a simple vertical scaling of the PSTH. For other electrode channels, however, the inclusion of additional units via re-thresholding with a more permissive threshold does change the spatial and temporal tuning by adding neurons with different tuning properties (Figure 2D). Interestingly, however, when we consider all electrode channels together and reduce the dimensionality of the data using PCA, we find the resulting low-dimensional projections are not sensitive to these individual channel-level changes. For both the randomly recombined units and the discarded spikes, the resulting neural population analyses replicate the original findings, including hypothesis tests evaluate with statistical criteria (e.g., p values), and the resulting low-dimensional neural population state trajectories closely resemble those estimated using only well-isolated neurons.

### Study 2: Neural population dynamics during reaching

We tested this analysis method on a second study, conducted by Churchland and colleagues, who discovered that neural population activity in motor cortex exhibits strong rotational dynamics during reaching Churchland et al. (2012); Pandarinath et al. (2015). The population rotational dynamics were revealed using jPCA, a dimensionality reduction algorithm that identifies 2D planes exhibiting rotational dynamics within the higher-dimensional neural state space. This study argues that motor cortex uses a set of oscillatory basis functions to construct the complex time-varying signals required to control muscles during a reach. Further statistical support for this observation has been provided provided by Elsayed & Cunningham (2017), demonstrating that the rotational dynamics are not a trivial consequence of smooth firing rates and applying dimensionality reduction algorithms to high-dimensional spaces.

The rotational dynamics described in Churchland et al. (2012) arise from limited-duration oscillatory patterns present in the firing rates of individual neurons, which poses a more challenging circumstance for testing whether multi-unit threshold crossings would reveal the same low-dimensional neural structure. A potential concern is that linearly combining the multiple units recording on a single channel would distort the observed population dynamics by “washing out” temporally-precise tuning features of individual units, since each of the constituent units in the multi-unit are not constrained to have similar relationships with behavioral parameters or time-varying patterns of activity. Thus, for this particular study, we might expect that combining oscillatory units with different periodicity or phases could decrease or eliminate rotatory dynamics at the level of the population.

We did not find this to be the case, and instead were able to recapitulate Churchland and colleagues’ findings using multi-unit threshold crossing data. Applying jPCA to the original spike sorted data yields a 2D jPCA plane which captures 23% of the variance in the neural data. Applying the same analysis to threshold crossing data yielded planes which capture 23%, 23%, and 21% of the variance in neural data when using voltage thresholds of −3.5, −4.0, and −4.5 × RMS, respectively. The key rotatory structure in the neural population data reported by Churchland and colleagues was also preserved and clearly present when using multi-unit threshold crossing data (Figure 3).

**Figure 3:**
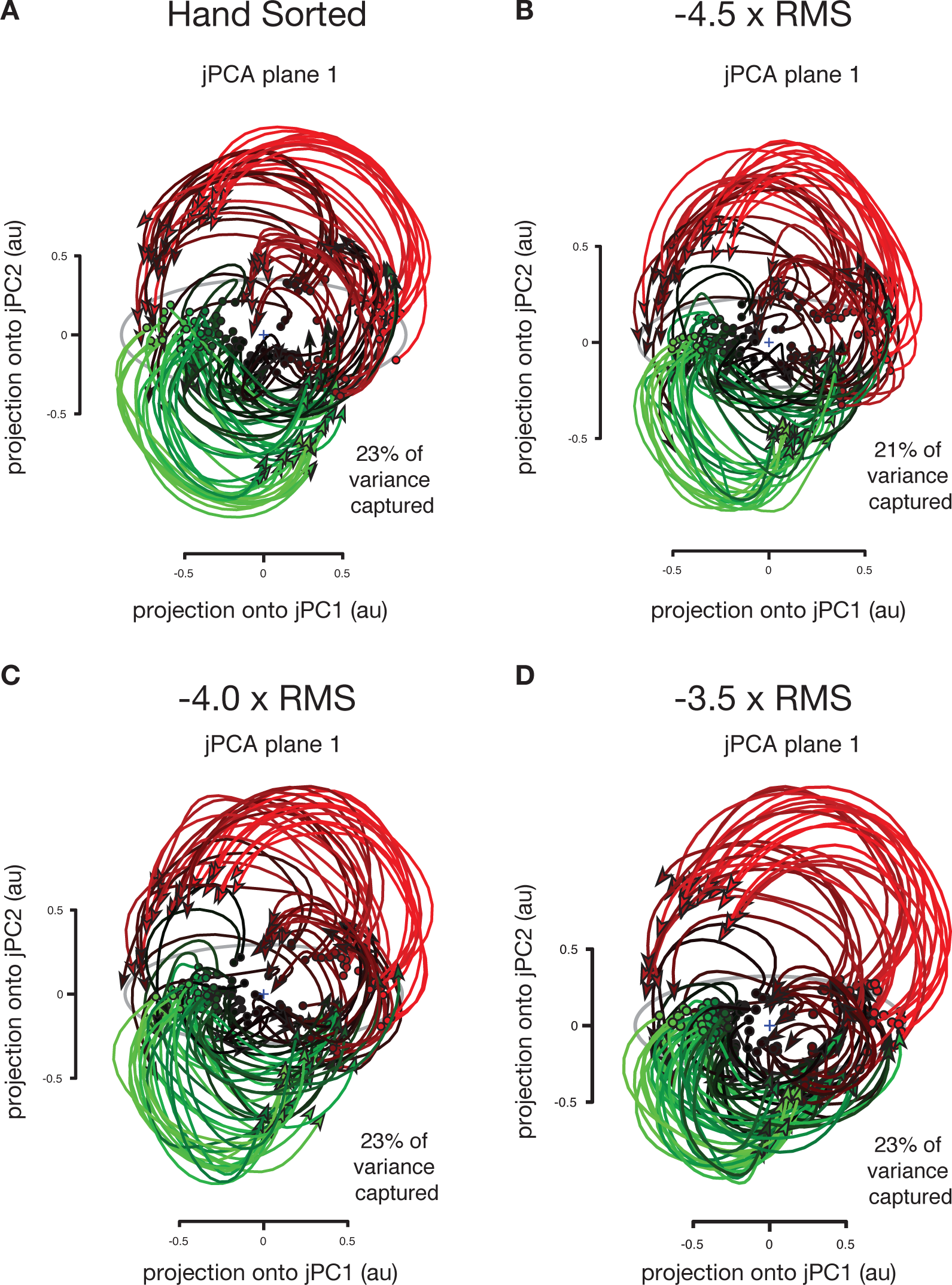
Replication of “Neural population dynamics during reaching” Churchland et al. (2012). Neural trajectories from 108 conditions including straight and curved reaches using… **(A)** Hand sorted units. **(B-D)** Multiunit activity using a voltage thresholded at −4.5, −4.0 and −3.5 × the root mean square (RMS) of the overall electrical recording. The total amount of variance captured in the top rotational plane as well as qualitative features of neural population state space trajectories is similar across sorted units and all three threshold crossing levels.

### Study 3: Cortical activity in the null space: permitting preparation without movement

The third and final study we replicated sought to understand how there can be large changes in PMd and M1 firing rates during a preparatory instructed delay period, without causing the circuit’s downstream muscle targets to move Kaufman et al. (2014). The mechanism the authors proposed is that the brain makes use of specific “output-null” dimensions in the neural state space (i.e., weighted combinations of firing rates of neurons that cancel out from the perspective of a downstream readout) to enable computation within a given circuit without influencing output targets. Other “output potent” neural dimensions do not cancel out, resulting in signals which do cause muscle activity. Kaufman and colleagues showed that neural population activity patterns before movement lay in the putative output-null subspace, consistent with this ‘output-null hypothesis’. Importantly, the output-null and output-potent dimensions were not separate sets of neurons (such as separate pools of delay-active neurons and movement-only neurons) Kaufman et al. (2014). Instead, these output-null and output-potent neural dimensions consisted of different weightings of the same neurons which were identified by contrasting the dominant modes within the lowdimensional population activity (found using PCA) between preparation and movement. Recently these dimensions have been shown to be truly orthogonal Elsayed et al. (2016).

Kaufman and colleagues’ findings represent a particularly strict and challenging test of the threshold crossing approach, as one might expect that combining multiple neurons might mix together output-potent and output-null dimensions. On the other hand, since these dimensions are weighted sums of many different neurons, then one might expect that that combining sums of neurons would provide similar results. Indeed, we find that with 192 threshold crossing channels (96 electrodes in PMd and 96 electrodes in M1), we observed the same distinction between output-potent and output-null neural dimensions as in the original study (Figure 4). This was quantified using the ratio of variance of neural activity in output-null to output-potent dimensions, found to be 3.84 (*p* = .048) using threshold crossings (−3.5×RMS), compared with tuning ratio 5.6 (*p* = .021) (Figure 4, dataset N20100812). In addition, we observed a larger transient component in both the output-null and output-potent response following target appearance, as shown in Figure 4C,D. Despite these differences, the multi-unit threshold crossing analysis of these data recapitulated the key finding that motor cortical firing patterns are largely restricted to output-null dimensions during movement preparation, which provides an explanation of how this activity is prevented from prematurely causing movements.

**Figure 4:**
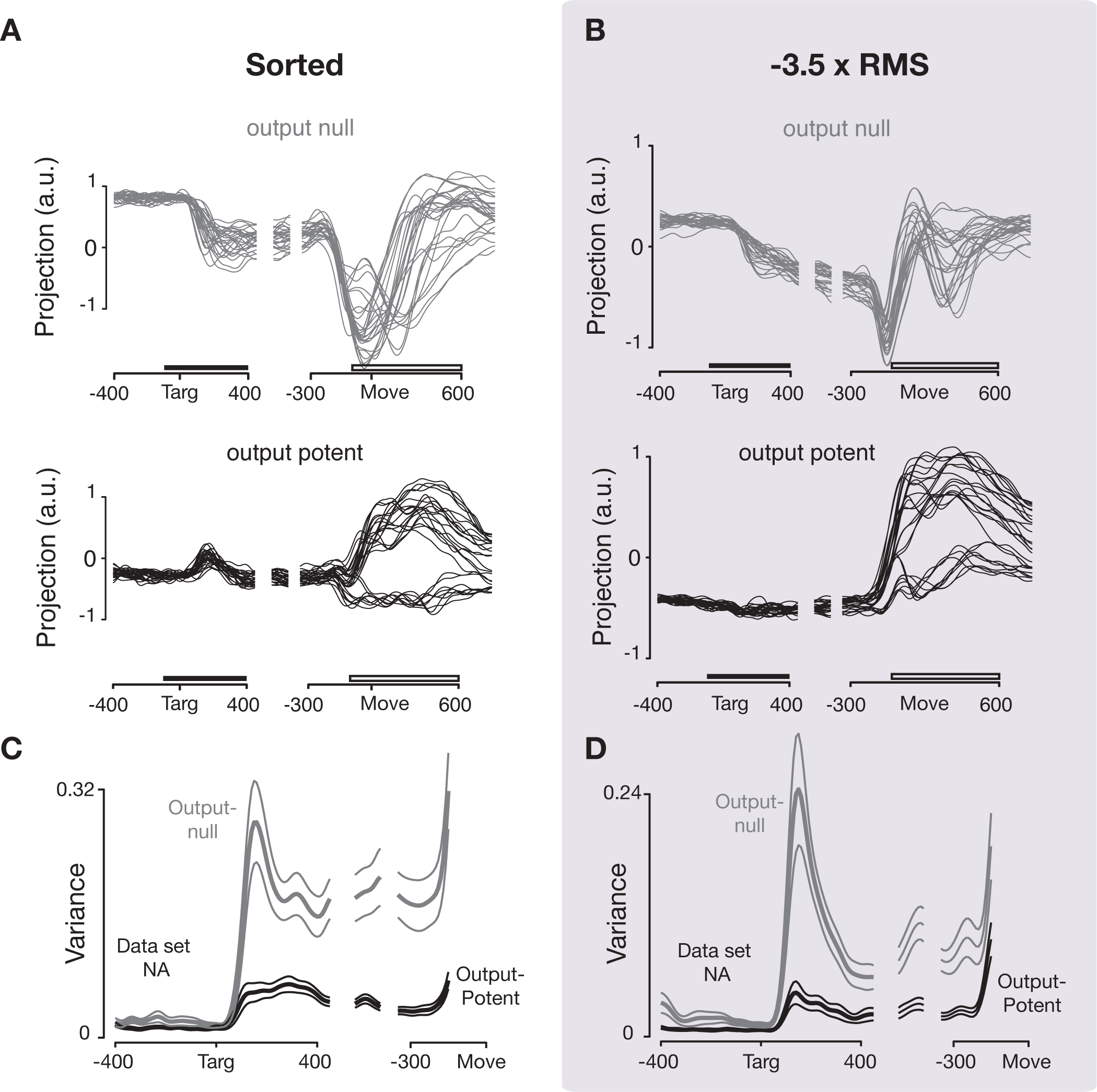
Replication of “Cortical activity in the null space: permitting preparation without movement” Kaufman et al. (2014). Comparison of output-null results from Kaufman et al. 2014 using sorted and thresholded data. **(A)** Neural activity in one output-null and output-potent dimension for one data set (NA), as in Figure 4A,B in Kaufman et al. (2014). Activity is trial-averaged, and each trace presents the neural activity for one condition. **(B)** Same as (A), computed using activity thresholded at −3.5 × RMS. **(C)** Tuning depth at each time point in output null and output potent dimensions, as in Figure 4C in Kaufman et al. (2014). **(D)** Same ac (C), computed using activity thresholded at − 3.5 × RMS.

Kaufman and colleagues’ study assessed whether preparatory activity occupies an orthogonal subspace to that comprising the movement activity. This analysis uses PCA to find a 6D space, then partitions that space into two 3D subspaces, an *output potent* and *output null* subspace. All neural activity, regardless of task-relevance, must exist in either of these two orthogonal subspaces, making this analysis intrinsically hungry for statistical power. The p-values calculated using the original study’s most conservative metric (which includes all neural data following target presentation, both initial transient and steady state) were 0.269 and 0.554 for thresholds of −4.0 × RMS and −4.5 × RMS, respectively (compared to X and Y using sorted neurons). The effect sizes are 2.006 and 1.926, respectively. The p-values calculated at steady state, however, were 0.042 and 0.119 respectively, and the variance tuning ratio plots are qualitatively quite similar. While these results are close, but do not quite both reach a p-value of 0.05 significance level, we speculate that the statistical power would grow significantly with additional channels of threshold crossing data as was used for the other array dataset in Kaufman et al. (2014). A germane independent validation of measuring the output-null hypothesis using threshold crossings (rather than sorted spikes) comes from our recent study which used BMI experiments to definitively show that the output-null mechanism isolates visuomotor feedback from prematurely affecting motor output Stavisky et al. (2017).

To summarize our experimental analysis results, we selected three recent studies based on the potential challenges and insights they could offer for this investigation of using multi-unit threshold crossings instead of sorted single unit activity. For all three previously published studies considered here, the key scientific advances have been recapitulated using threshold crossings instead of individual neurons. While this outcome need not be the case for all studies, the fact that it worked for these three different studies, which span a variety of datasets, questions, and analysis techniques, suggests that it will generalize to many other population-level analyses of neural activity. This could enable many new scientific analyses, and we anticipate its relevance and importance to continue to grow as the number of recorded neurons in experiments increases.

### A random projection theory of recovering neural population dynamics using multi-unit threshold crossings

We observe that the low-dimensional neural population dynamics reported here using threshold crossings are very similar to those reported in previous studies that relied on single neurons isolated using spike-sorting methods. Given that combining action potentials from several neurons on each electrode channel discards some information, this result may seem surprising. Why does discarding information have such a small effect on the estimated dynamics? Here we use the theory of random projections to provide a quantitative explanation. Central to this explanation is the concept of a manifold, or a smooth, low dimensional surface containing the data. This explanation reveals that when a manifold embedded in a high dimensional space is randomly projected to a lower dimensional space, then the underlying geometry of the manifold will incur very little distortion under the following conditions: the manifold itself is simple (i.e., smooth, with limited volume and curvature) and the number of projections is sufficiently large with respect to the dimensionality of the manifold. While true theoretically, the key practical question is, does this theory guarantee accurate recovery of neural population dynamics under experimentally and physiologically relevant conditions?

To apply random projection theory to our data, we must first define the high-dimensional space and the low-dimensional manifold. In this neuroscientific application, the high dimensional space has one axis per neuron, with the coordinate on that axis corresponding to the firing rate of that neuron. Thus, the space’s full dimensionality is equal to the total number of neurons in the relevant brain area, which is much larger than the total number of neurons from which we are able to record. However, in many brain areas, particularly within the motor system, most neurons’ trial-averaged neural activity patterns vary smoothly across both time and behavioral-task conditions and exhibit consistent correlation structure between neurons. Thus, as one traces out time and conditions, the resultant set of covarying neural activity patterns constitutes the embedded manifold in random projection theory Gao & Ganguli (2015) (i.e., blue curve in top-left of Figure 5A). Empirically, we find that the dimensionality of neural activity is far lower than the number of recorded neurons. This is likely due in large part to the structure of connections within the network, but may also be in part due to the fact that the experimental conditions do not fully span the set of possible behaviors.

**Figure 5:**
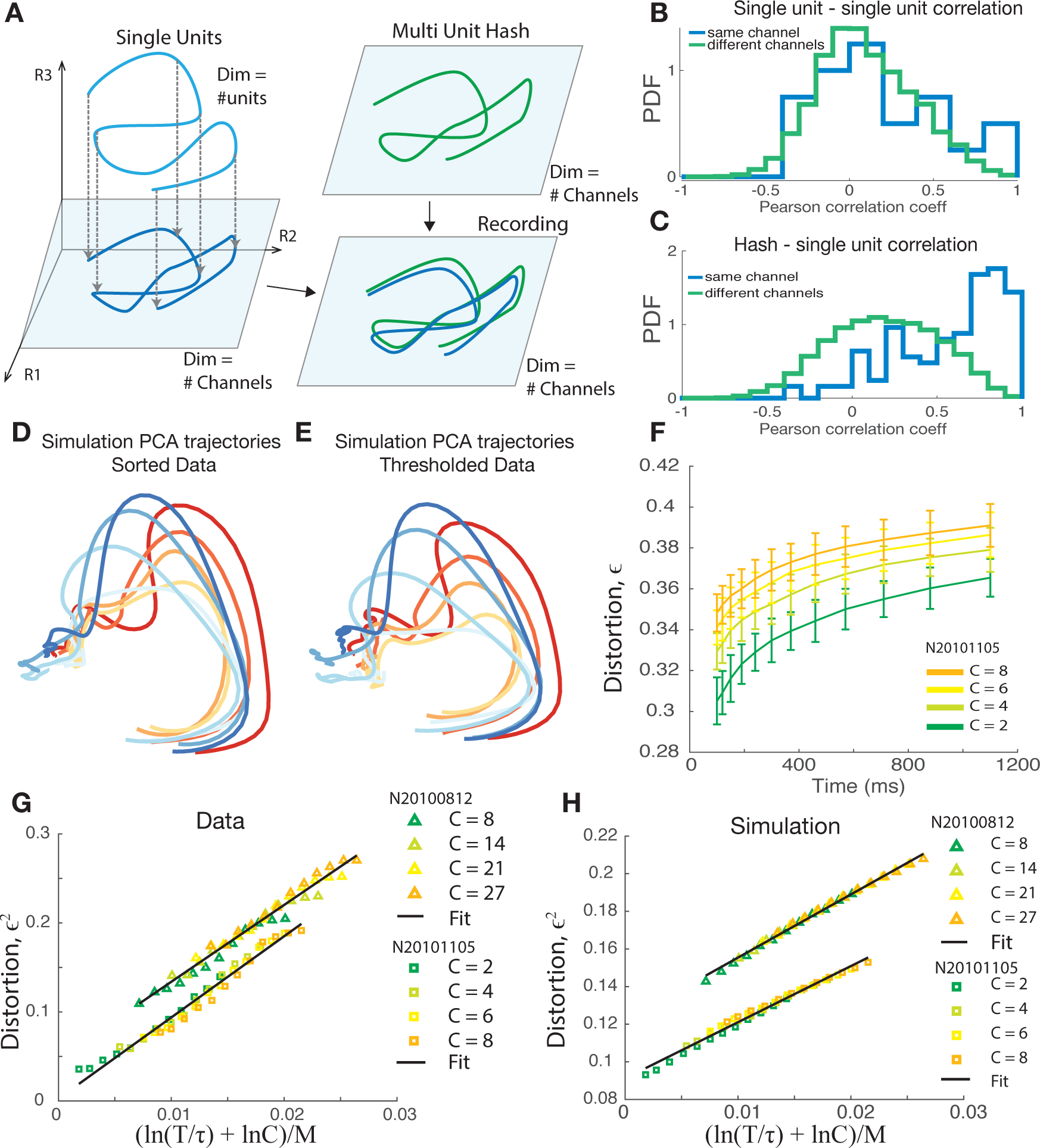
Random projection theory supports not spike sorting. **(A)** Schematic depiction of a projection of a trajectory through high-D firing rate space defined by single units, projected onto a subspace defined by the number of recording channels. A small amount of information is lost by combining units on each channel. Adding the contribution of multi-unit hash may introduced additional distortion to the estimated neural trajectories, though in practice this appears to be small. **(B)** Pearson correlation coefficient between single units on the same recording channel (blue) and different channels (green) for dataset N20101105. **(C)** Hash has a larger Pearson correlation coefficient (*p* < 0.05) with single units on the same channel (blue) than from other channels (green). Same dataset as (B). **(D)** PCA trajectories sampled from simulated random Gaussian manifolds were measured from (simulated) single neuron activities. Manifold mean and covariance were matched to those of neural activity from dataset N20101105 spike sorted data. **(E)** the same as (D) for threshold crossings data. **(F)** The maximum distortion of random onedimensional manifolds under random projections of *N* = 125 neurons. The length of the manifolds, *T*(*sec*), and the number of manifolds, *C*, are varied with fixed correlation length, τ = 14.1. The 95th percentile of the distortions under 100 random projections is plotted (mean ± standard deviation for 50 repetitions). This collapses into a simple linear relationship when viewed as a function of ln(*CT*/τ), plotted for data **(G)** and for simulated random Gaussian manifolds **(H)**.

Empirically, we find that activity in motor cortex is much lower dimensional than the number of neurons, limiting the manifold volume. The manifold also has limited curvature, precisely because neural activity patterns vary smoothly across time and conditions. Thus this manifold satisfies the condition of simplicity posited in random projection theory.

Finally, the mapping from single neuron firing rates to threshold crossing activity constitutes a random projection itself: each electrode’s activity is a weighted linear combination of a small number of isolated neurons plus any additional hash that passes the threshold. More specifically, threshold crossings are not only a linear weighting but an equal weighting of each neuron’s action potentials (i.e., multi-unit threshold crossing rate = neuron 1’s rate + neuron 2’s rate + …). Thus, the low dimensional space in random projection theory has dimensionality equal to the number of electrodes. Within this low-dimensional space of electrodes (which itself is a subset of the high dimensional space of neurons), neural activity traces out a simple manifold (Figure 5A) due to the even lower-dimensional latent dimensionality of the activity.

This raises the critical question: how much is the geometry of the manifold distorted when combining single-neuron firing rates within each electrode? To help understand this question, we generated random manifolds of neural activity using simulated single units and multi-unit firing rates, and estimated the dynamics of these random neural manifolds. In generating these synthetic datasets, we incorporated the average correlation structure between single neurons (Figure 5B) and between single units and multi-unit threshold crossings on a given channel (Figure 5C). Neural population state-space trajectories obtained from simulated single neurons (Figure 5D) closely match those obtained from simulated threshold crossings (Figure 5E). These results are consistent with and reinforce the previous sections’ comparisons of single unit and threshold crossing neural dynamics in real neural data.

Random projection theory enables us to go beyond this qualitative view and derive a quantitative theory of how the geometric distortion between threshold crossings and single-neuron neural population state-space trajectories depend on their underlying complexity and on the number of measurement channels Donoho (2006); Baraniuk & Wakin (2009); Ganguli & Sompolinsky (2012); Advani et al. (2013). Here we define the distortion as the worst case fractional error in distances between all pairs of points on all trajectories, measured in the space of electrodes, relative to the space of single neurons (see Methods, and also Lahiri and colleagues for a detailed discussion Lahiri et al. (2016)).

This distortion will depend on the volume and curvature of the trajectories. In the simple case where neural population dynamics consists of *C* different neural trajectories each of trial duration *T*, the volume can be taken to be proportional to *CT*. In turn, the curvature is related to the inverse of the temporal auto-correlation length τ of the neural trajectories Clarkson (2008); Baraniuk & Wakin (2009); Verma (2011); Gao & Ganguli (2015); Lahiri et al. (2016). Intuitively, τ (see Methods) measures how long one must wait before a neural trajectory curves appreciably, so that small (large) τ indicates high (low) curvature.

We can parametrically vary the complexity of neural population dynamics by analyzing subsets of neural trajectories of different durations *T* and subsets of conditions of different sizes *C*. In doing so for real data, we find that the distortion grows with both *T* and *C* (Figure 5F). However, random projection theory predicts a very striking and specific scaling relation between the squared distortion ϵ^2^ and the parameters *T*, *C*, and number of channels *M*. In particular it predicts that ϵ^2^ scales linearly with 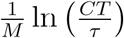. Thus, distortion is predicted to be directly proportional to the logarithm of the product of manifold volume and curvature (i.e. 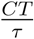), and inversely proportional to the number of recording channels M. This predicted scaling relation is seen both in neural data (Figure 5G) and in simulations where the data was generated so as to match the overall average statistics of the recorded data (Figure 5H).

We note, however, that the distortion relationships found in the two different datasets (and the simulated data generated based on their statistics) form two different lines in (Figure 5G,F). A key reason for the discrepancy is that these analyses deal not with deterministic random projections, but rather noisy random projections, in which the presence of hash introduces both signal and additional noise, or activity which is uncorrelated to the single neurons on a channel. A natural measure of signal to noise is the ratio of the variance of single unit activity, to the variance of that part of the hash that is not correlated with the single units. By fitting a generative model to the neural data, we can estimate this SNR (see Methods). We found it to be 1.9 for dataset N20101105 in Ames et al. (2014) and 1.3 for dataset N20100812 in Kaufman et al. (2014). Thus, as expected, both the recorded and simulated data incur higher distortion at fixed complexity 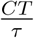 if the SNR is lower.

How does the SNR more generally impact distortion? While there is as yet no general theory of how noisy random projections distort the geometry of smooth manifolds, we empirically find the scaling relation to be 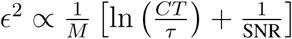, as verified in simulations in Fig 5H. Thus this novel scaling relationship provides quantitative theoretical guidance for when we can expect to accurately recover neural population dynamics without spike sorting. Intuitively, analyzing threshold crossings makes sense when (1) neural trajectories are not too long (small *T*, e.g.: less than a few seconds), (2) not too curved (large τ), (3) not too many in number (small *C*), (4) the additional multi-unit hash has small variance relative to that of single-units on the same electrode (non-negligible SNR), and (5) we have enough electrodes (large *M*). Under these conditions, neural dynamics can be accurately inferred using multi-unit threshold crossings. All of these conditions are satisfied in the datasets considered here, and are likely to be satisfied in many more. Future studies of recordings from other brain regions, in other species, and across a range of conditions (1-5 above) will be needed to further establish the generality of these experimental, analytical and theoretical findings.

## Discussion

We investigated whether multi-unit threshold crossings can be used instead of isolated single neurons for questions involving neural population dynamics. We believe that this question is necessary and timely, as neuroscientists and neuroengineers are currently engaged in a massive and expensive scale-up in the number of electrodes used to make measurements. Moreover, the field is facing the question of the type of neural data (e.g., single neurons, multi-units, local field potentials) that will be important to record when thousands to millions of electrodes are available. Some of these questions are already starting to be confronted in the related but distinct endeavor of large-scale neural imaging using genetically-encoded calcium indicators.

We report here both an empirical validation and a theoretical justification which together argue that many of the same scientific conclusions can be made about motor cortical population activity without spike sorting. We believe that these findings are broadly useful for several reasons. First, the process of spike sorting is both time consuming and inexact, with significant variability between experts Wood & Black (2008). For a typical experiment composed of several hours of neural recordings, an expert human sorter may spend several hours to manually sort spikes on a single 100-channel electrode array (e.g., “Utah array” Maynard et al. (1997)). New high-channel-count recording technologies are becoming available that will enable recording from thousands to millions of channels simultaneously. A data set composed of 1,000 channels could take over 100 hours to hand sort, with no ground truth available to validate results. As reported here, this effort has the potential to yield little if any impact on the resulting scientific insight.

Second, in real-world experimental conditions, chronically implanted multi-electrode arrays in animal models or in humans often feature many channels with neural activity that cannot be isolated into single neurons (e.g., Pandarinath et al. (2015)). Ignoring these channels throws out potentially meaningful task-related information, possibly weakening statistical power and subsequent scientific conclusions. Insisting on spike sorting in these situations would fail to capitalize on valuable experimental and clinical opportunities. Including such electrodes will enable analyses of vastly larger datasets from neurons in more brain areas; permit more efficient use of experimenter time by avoiding time-consuming manual spike sorting; and reduce the number of research animals and clinical participants needed, by making use of electrode sensors with less pristine recordings due to device age or random variability.

Third, these results support the design of novel classes of sensors for both scientific experimentation and BMI clinical trials, as described in more detail below.

Finally, and from a broader scientific perspective, this study may help increase scientific reproduceability and reduce the number of animals or clinical trial participants required for a given study. Regarding reproduceability of findings, removing the often subjective step of spike sorting, which typically includes only vague descriptions of how spike sorting was accomplished, ought to increase the ability of multiple labs to replicate results from the same data set or to replicate findings as part of subsequent studies. Regarding reducing the number of subjects needed, by quantitatively and theoretically grounding the approach of using threshold crossings, large numbers of unused existing data sets can now be brought to bear on new hypotheses. Data sets collected using older electrode technologies, or where biological conditions prevent detection of well isolated cells, are likely appropriate for answering new questions without spike sorting, as opposed to the more stringent spike sorting step, and thereby potentially reduce the number of new subjects required.

We stress that this method is not appropriate for scientific questions that seek to investigate the properties of individual neurons (i.e.: not for questions regarding stimulus selectivity, single cell tuning properties, etc). For such studies, we must either rely on closely clustered electrodes and automated sorting methods or restrict the number of neurons to a feasible hand-sortable quantity.

### Necessary conditions for using this paradigm

The Johnson-Lindenstrauss lemma shows that manifold estimation error increases as the complexity of the underlying manifold increases Johnson & Lindenstrauss (1984). The studies replicated here have all focused on neural recordings from PMd and M1, where roughly 10-15 dimensions capture 90% of the variability of trial-averaged neural population activity when monkeys perform a simple 2D reaching task (e.g., Yu et al. (2009)). Recording from one or two Utah arrays (i.e., 96 or 192 electrodes) provides sufficiently redundant sampling. In this regime, we robustly find that we can replicate scientific hypotheses using electrical threshold crossings in place of well isolated single units.

The observation of low-dimensional neural population dynamics is well established in the motor system, but may not be true for all brain areas. In other brain areas, particularly input-driven sensory areas, the dimensionality of neural activity may be significantly higher or scale rapidly with the complexity of a sensory stimulus, resulting in more complex activity manifolds Cowley et al. (2016). Under these other conditions, it remains to be determined how many independent random projections (multi-unit recording channels) would be required to accurately recover the underlying structure of neural activity.

### Implications for neural recording sensor design

These findings unlock the potential for developing both acute and chronic multielectrode arrays which feature many thousands to millions of electrode channels at the expense of discarding information necessary to sort individual neurons.

Virtually all existing acute and chronic electrode arrays are designed with the ability to record broadband analog data from each channel (e.g., digitizing 30,000 samples per second). Relaxing these constraints reduces the storage and processing requirements for acute experiments and enables low power, high-channel count devices for clinical applications, where size, power, and communication bandwidth requirements constrain the number of channels. Regarding acute experiments, saving only the spike times for multi-unit threshold crossings reduces the total data volume between 4 and 5 orders of magnitude relative to saving broadband data. This is especially relevant for clinical applications, where wireless neural data communication is necessary. Radios consume considerable power at high bandwidth, which increases with sampling rate and bit depth. This work argues in favor of developing chronically-implantable electrode arrays with integrated electronics and wireless-data transmission, which optimize for low power consumption and communication bandwidth in place of spike sorting, enabling dramatically higher total channel count given a space and power budget (e.g.: Chestek et al. (2009); O’Driscoll et al. (2011).

Second, for any fixed number of electrodes to be distributed throughout some volume of tissue (e.g., across the surface of the cortex, in depth through cortex, as well as in deeper structures), there exists a trade-off between the goals of spike sorting, which benefits from closer electrode spacing, and recording from a larger number of neurons distributed through a larger volume. For example, the newly-developed Neuropixels probes enable a user to select a subset of 384 active channels from a total of 966 along a single-shank silicon probe. By selecting a sparse subset of the active channels, a user can elect to trade off additional tissue coverage for spike sorting quality at the outset of an experiment Jun et al. (2017b).

This study reinforces the idea that selecting an appropriate sensor depends on both the scientific goals of an experiment and the structure of activity in the recorded brain region. In brain areas whose neural population activity is governed by low-dimensional dynamics, a sensor that prioritizes quantity of independent neural channels over quality of unit isolation may be appropriate.

## Conclusions

The experimental neuroscience paradigm proposed here, using threshold crossings, applies to neural population-level analyses. It is not applicable in cases where one wishes to make statements about the properties (e.g. stimulus selectivity) of individual neurons. We nonetheless anticipate that this approach is broadly applicable to systems neurophysiology and is relevant not only to the analysis of experimental data, but also to the design and use of new chronic electrode arrays and acute multi-site recording probes. Using threshold crossings in lieu of spike sorted units will likely become increasingly important and enabling for population analyses in order to address growing dataset sizes and to enable next-generation brain-machine interfaces with considerably greater capabilities, performance and robustness. The present work demonstrates that this method is theoretically justified, empirically supported, and simple to use.

## Methods

Details of the experimental procedures for the three previous studies whose data was reanalyzed here using threshold crossings, including these studies’ behavioral analyses, neural recordings, and neural analyses, have been detailed previously in Ames et al. (2014); Churchland et al. (2012); Kaufman et al. (2014). In all three of those experiments, monkeys performed a delayed point-to-point reaching task to targets presented on a touch screen while neural data was recorded from two Utah arrays (Blackrock Microsystems, USA) placed in PMd and M1 of the contralateral hemisphere. All surgical and animal care procedures were performed in accordance with National Institutes of Health guidelines and were approved by the Stanford University Institutional Animal Care and Use Committee.

### Data Preprocessing

In all three sets of experiments, raw neural recordings (voltage measurements taken at 30,000 Hz) were first high-pass filtered above 250 Hz (4-pole, “spikes medium” setting) by the Blackrock neural recording system. Then, voltage snippets around measurements that crossed a threshold at −3.5 × root mean square voltage level for a given channel were saved. Spike sorting for the original studies was performed by hand, based on grouping spikes by similarity of waveform (i.e., assigning them to putative single neurons). For the present study’s re-analyses, ‘re-thresholded’ datasets were constructed by calculating new voltage threshold levels for each channel, and retaining the subset of original waveforms that also crossed the more conservative thresholds at −4.0 and −4.5 × RMS. Threshold crossings, regardless of waveform shape, were treated as spike events and processed further into temporally-smoothed, trial-averaged PSTHs prior to subsequent analyses such as PCA Ames et al. (2014); Kaufman et al. (2014) or jPCA Churchland et al. (2012).

The result obtained by Kaufman and colleagues is particularly hungry for statistical power, as it simply involves a binary partition of a 6D space, and looks for differences in activity at different time points in the two subspaces. For this study, after re-thresholding data, noisy and/or very weakly modulated units were removed using an SNR threshold. For this context, SNR as a unit’s modulation depth (i.e., maximum minus minimum firing rate) across all conditions and time, divided by the peak of that unit’s standard deviation across all conditions and time. Sweeping the SNR rejection threshold between 0 and 1.2 results in rejecting 0 to 8.8% of units, respectively (-3.5 rms dataset). The calculated p-value varies between a maximum of 0.065 (no unit filtering) and a minimum of 0.0475 across this range.

### Definition of the Distortion of Manifolds under a Projection

Let r be a high-dimensional vector of single neuron firing rates at an instant of time. Let e be the corresponding thresholded, temporally smoothed electrode activities at the same time. They are related via a noisy projection e = Ar + h where the elements of the matrix A indicate how much each neuron contributes to each electrode, and the noise vector h indicates the additional contribution of hash that passes threshold. The geometric distortion of a single electrode pattern e, relative to a firing rate pattern r is defined as

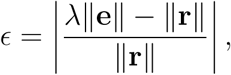

where λ is a scale factor, used to remove the effect of multiplication of all vectors by a single scalar. Here ||·|| denotes the length of a vector, and so the distortion is the fractional change in length up to a scale λ.

The distortion of a set of manifolds is computed by finding the maximum distortion of the vectors between all pairs of points on any of the manifolds, including those between different manifolds. The scale factor λ is chosen to minimize the maximum distortion of the manifold set, thereby correcting for trivial overall changes in scale induced by the projection. Thus the distortion of a set of manifolds is the worst case fractional error in pairwise distances, up to an overall scale.

### A generative model for neural manifolds and their projections

To quantitatively test how well the framework of random projections of smooth manifolds matches neural data, we developed a generative model of neural state space dynamics in single neuron firing rate space, its projection to electrodes, and the addition of hash. We modeled the neural population dynamics themselves as a random smooth manifold. We generated 50 different such manifolds and their noisy projections by sampling *T* × *C* × (*N* + *M*) numbers (firing rates) from a multivariate Gaussian distribution, with mean and covariance chosen to have the same statistics as in the data, where *T* is the number of time points in a neural trajectory, *C* is the number of conditions, *N* is the number of single units and *M* is the number of recording channels (192 in this case). The multivariate Gaussian distribution we used corresponds to the following generative model for the PSTHs of single units, *r_n_*, multi-unit hash, *h_μ_*, and channels, *e_μ_*:

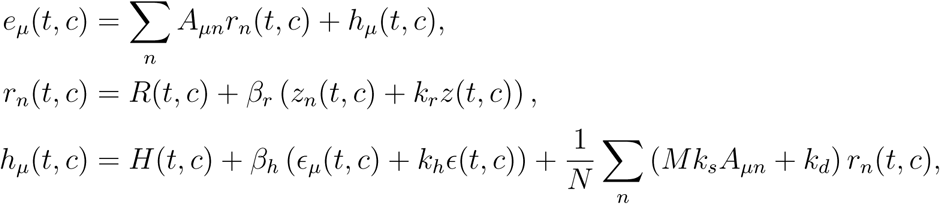

where *z_n_*(*t*,*c*), *z*(*t*,*c*), *ϵ_μ_*(*t*,*c*), and *ϵ*(*t*,*c*) are independent Gaussian random processes whose autocovariance across time and conditions were set equal to corresponding quantity computed from data, normalized to have unit total variance. The functions *R*(*t*,*c*), *H*(*t*,*c*), and the constants *β_r_*, *k_r_*, *β_h_*, *k_h_*, *k_s_*, *k_d_*, were fit to the data by setting the means, standard deviations and correlation coefficients of *r_n_* and *h_μ_* in the model equal to the mean of the corresponding quantity in the data. The constant *k_r_* accounts for correlations between single units, *k_h_* for correlations between hashes, *k_d_* for correlations between hash and single units, and *k_s_* for the stronger correlations between hash and single units on the same channel.

In this generative model it is possible to separate the simulated unsorted activity, *e_μ_*, into noise (the terms proportional to *β_h_*) and signal (everything else). This allows us to calculate a signal-to-noise ratio for each dataset after fitting the model parameters, defined as the ratio of the variances of signal and noise.

For each manifold set, 100 random projections were sampled, which were *M* × (*N* + *M*) matrices of ones and zeros, where each of the *N* single units contributes to one channel chosen randomly from *M* of them, and each of the *M* hashes contributes to its corresponding channel.

For each random projection and random manifold set, the distortion of the manifold set was computed as described above. The 95th percentile of the distortions under the 100 projections was recorded. The mean and standard deviation of these distortion percentiles was taken over the 50 manifold sets.

### Distortion of PSTHs

The PSTHs of the spike sorted data were taken to be the set of unprojected manifolds and the thresholded data was taken to be the set of projected manifolds. The distortion of the manifolds was then computed as described above.

### Correlations of PSTHs

The covariance matrix across time and conditions was computed from spike sorted data by treating each neuron as a sample. The normalized covariance matrix was computed by dividing the covariance matrix by the total variance across time and conditions. To compute the correlation time, this covariance was then averaged for pairs of times with the same separation. This covariance function was then fit to a sum of two Gaussians centered at zero. The correlation time τ was then computed by matching the value and second derivative at zero to that of a single Gaussian. The correlation coefficients and standard deviations of single units and hash were computed from spike sorted data by treating each time point and condition as a sample.

## Acknowledgements

We thank M. Risch, M. Wechsler, J. Aguayo, C. Sherman, E. Morgan, for surgical assistance and veterinary care; B. Davis, and E. Castaneda for administrative support; and B. Oskotsky for information technology support. This work was supported by the NIH NRSA 1F31NS089376-01 (E.M.T), Stanford Graduate Fellowship (E.M.T, C.K.A.), NSF GRFP (S.D.S., M.T.K.), the NSF IGERT 0734683 (E.M.T., S.D.S.), the Christopher and Dana Reeve Foundation (S.I.R. and K.V.S.), and the following to K.V.S.: Burroughs Wellcome Fund Career Awards in the Biomedical Sciences, DARPA REPAIR grant N66001-10-C-2010 and NeuroFAST grant W911NF-14-2-0013, NINDS grant R01NS076460, NICHD grant 8DP1HD075623-04, NIMH grant 5R01MH09964703, and the Simons Foundation. K.V.S. is a consultant to Neuralink Inc. and on the Scientific Advisory Boards of Cog-nescent and Heal; these entities in no way influenced or supported this work. S.G. and S.L are supported by the Burroughs Wellcome, Sloan, Simons, McKnight, and James S. McDonell foundations, and the Office of Naval Research.

## Author Contributions

E.M.T. and S.D.S. performed the analyses and wrote the manuscript. C.K.A. and M.T.K. provided data and analysis code. S.L. and S.G. provided theory work and simulation results. S.I.R. assisted with animal surgeries. K.V.S. provided extensive feedback.

## Supplemental Figures

**Figure S1:**
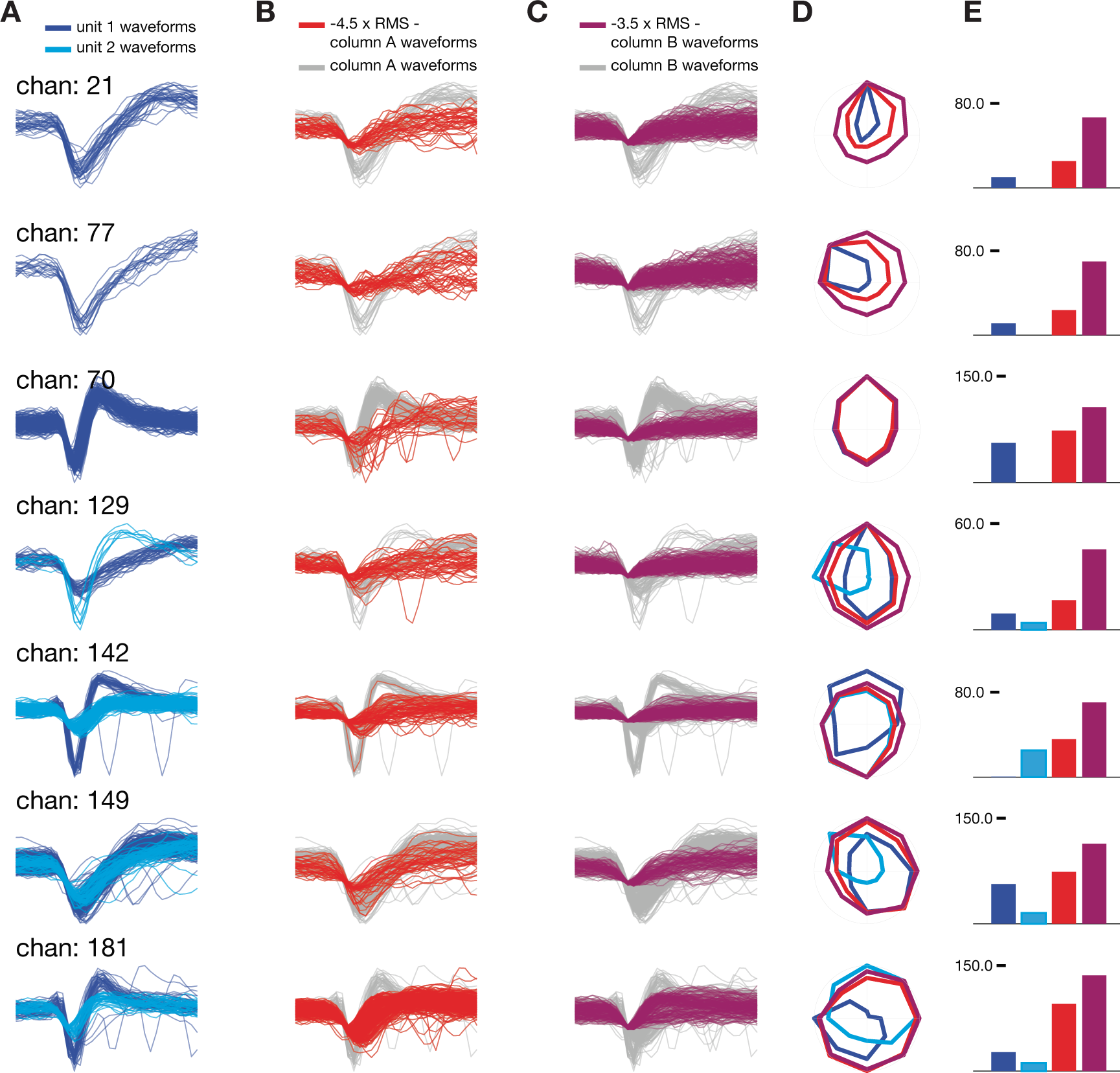
Waveforms and tuning of single and thresholded units. **(A)** Hand-isolated waveforms from channels with either one or two units. **(B)** All waveforms detected using signal threshold −4.5 × RMS. Additional waveforms not already present in (A) are highlighted in red. **(C)** All waveforms detected using signal threshold −3.5 × RMS. Additional waveforms added not already present in (B) are shown in purple. **(D)** Firing rate tuning curves during reaches to eight radially spaced targets for the three thresholding levels shown in A-C. Consistent with expectations, more permissive thresholds broaden tuning curves but generally preserve peak tuning direction. **(E)** Firing rate for peak tuning direction for each threshold level A-C. More permissive thresholds result in higher firing rates.

**Figure S2:**
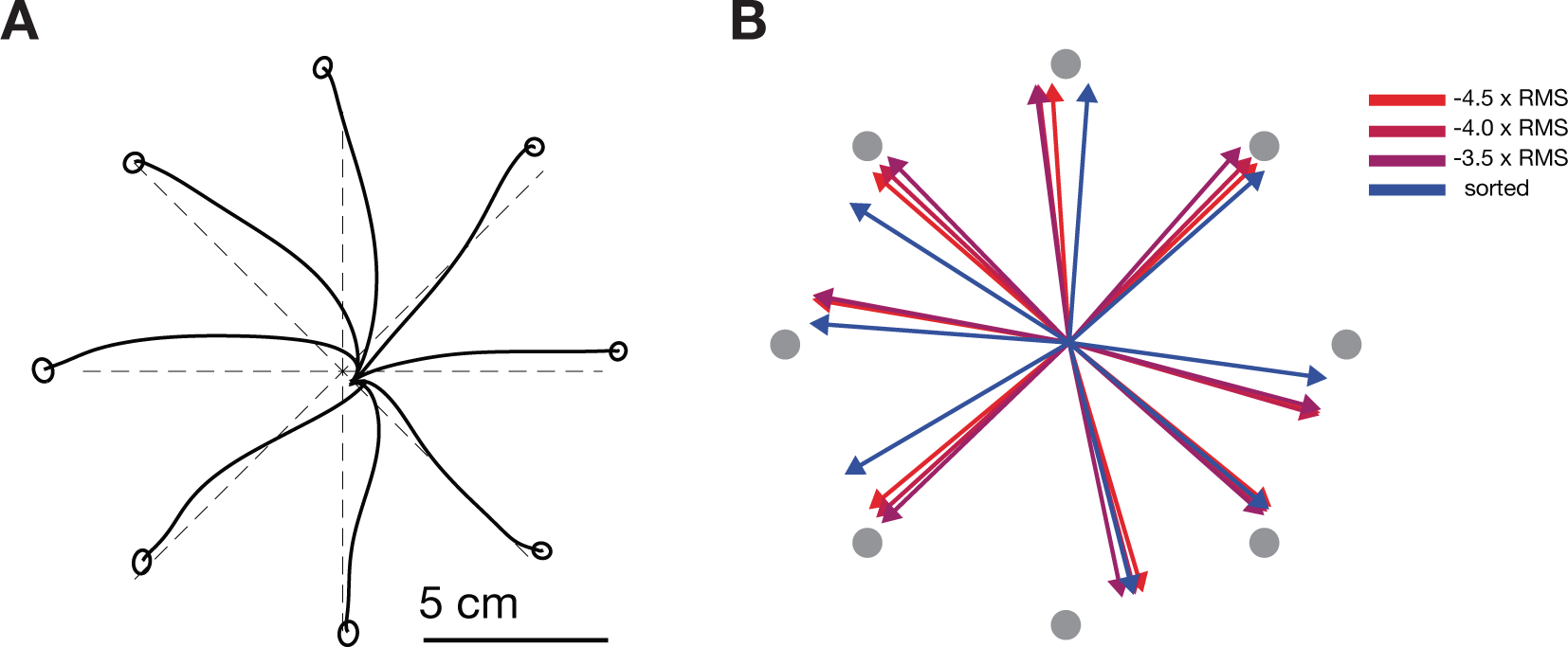
Waveforms and tuning of single and thresholded units. **(A)** Hand-isolated waveforms from channels with either one or two units. **(B)** All waveforms detected using signal threshold −4.5 × RMS. Additional waveforms not already present in (A) are highlighted in red. **(C)** All waveforms detected using signal threshold −3.5 × RMS. Additional waveforms added not already present in (B) are shown in purple. **(D)** Firing rate tuning curves during reaches to eight radially spaced targets for the three thresholding levels shown in A-C. Consistent with expectations, more permissive thresholds broaden tuning curves but generally preserve peak tuning direction. **(E)** Firing rate for peak tuning direction for each threshold level A-C. More permissive thresholds result in higher firing rates.

**Figure S3:**
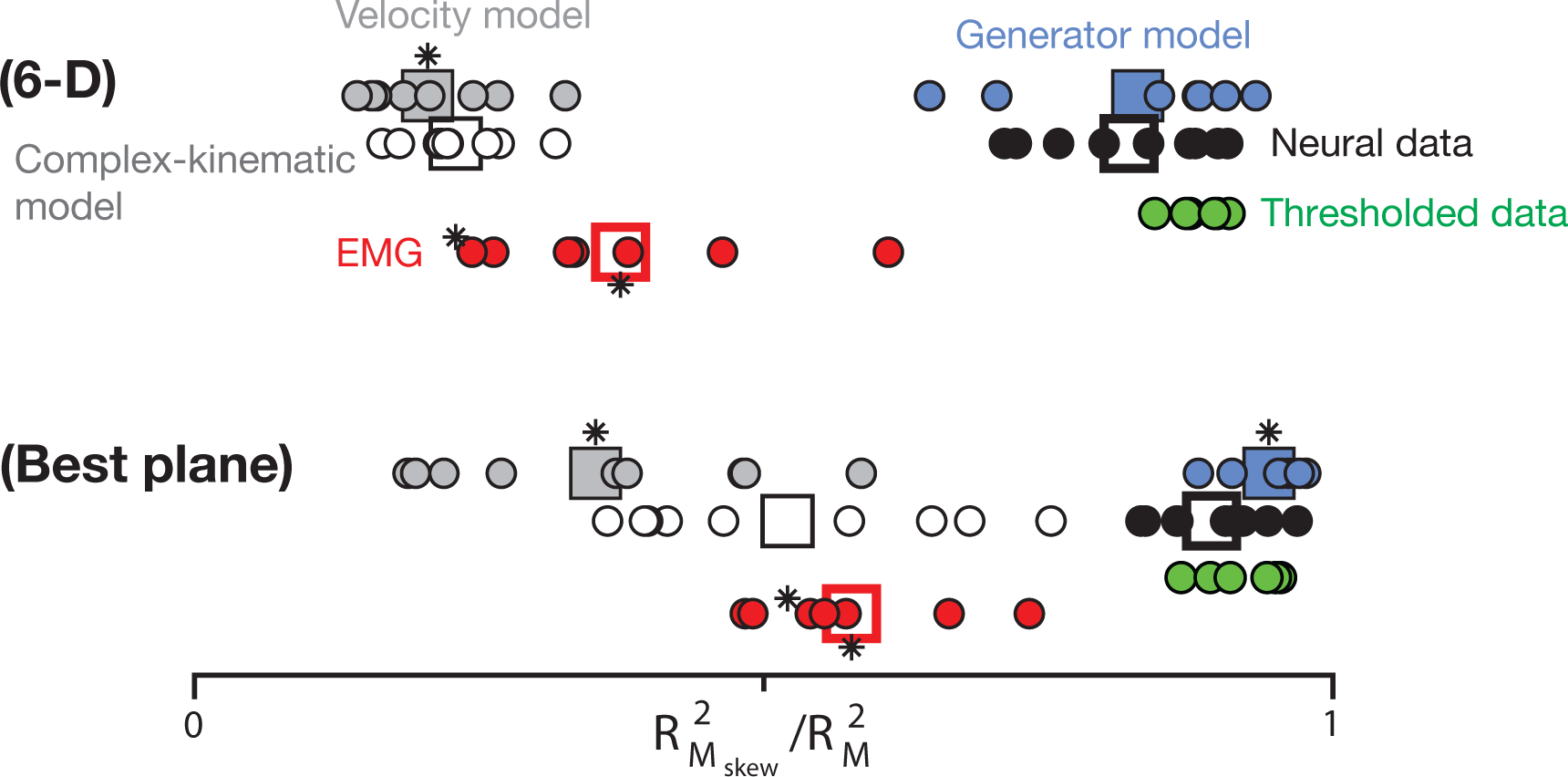
Consistency of rotational dynamics of thresholded data. Figure reproduced from Churchland et al. (2012), Figure 6, with additional data points for the re-thresholded data sets (green markers), illustrating that the ratio 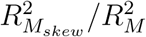 for thresholded datasets is consistent with hand sorted neural data.

